# Exploring differences across pangenome-graph representations using *Escherichia coli* O157:H7 as a model

**DOI:** 10.64898/2026.02.25.707890

**Authors:** Peilin Liu, Kaixin Hu, Lapo Mughini-Gras, Aldert L. Zomer, Michael S. M. Brouwer, Timothy J. Dallman, Julian A. Paganini

**Affiliations:** Faculty of Veterinary Medicine, Institute for Risk Assessment Sciences (IRAS), Utrecht University, 3584 CL Utrecht, the Netherlands; Centre for Infectious Disease Control, National Institute for Public Health and the Environment (RIVM), Bilthoven, Netherlands; Faculty of Veterinary Medicine, Department of Infectious Diseases and Immunology, Utrecht University, Utrecht, The Netherlands

## Abstract

Pangenome graphs are increasingly used to represent population-scale bacterial diversity, yet construction methods span fundamentally different representation paradigms whose outputs and sensitivities to assembly quality remain poorly quantified. We systematically reviewed microbial pangenome graph tools and benchmarked six representative methods spanning gene-cluster, compacted coloured de Bruijn graph (ccDBG), multiple sequence alignment, and hybrid approaches. Using a repeat-rich *Escherichia coli* O157:H7 dataset with complete genomes and matched short-read data, we constructed graphs from identical inputs and observed orders-of-magnitude differences in graph size and fragmentation, indicating that global topology is driven by representation strategy. Varying completeness composition revealed that assembly fragmentation is a first-order determinant of graph structure: gene-cluster graphs contracted as draft assemblies replaced complete genomes, whereas unitig graphs expanded, with distinct degree–prevalence fingerprints across tools. Computational cost mirrored these shifts and depended strongly on completeness composition, including a pronounced runtime penalty for one ccDBG implementation on all-draft inputs. Finally, analysis of Shiga toxin loci showed that pangenome-level reconciliation does not reliably correct assembly artefacts at challenging multi-copy genes and that performance varies by locus. Together, these findings show that pangenome graphs are representation-dependent models of bacterial diversity, and that assembly completeness is a primary determinant of their topology, scalability, and locus-level accuracy.

**Authors Summary:** Bacterial populations are often described using “pangenome graphs,” which aim to capture all genetic variation across many genomes in a single structure. However, different tools build these graphs in fundamentally different ways, and little is known about how those differences affect the results. In this study, we systematically compared several widely used approaches using a clinically important strain of *Escherichia coli* that is rich in repeated and mobile DNA. We found that the size, connectivity, and overall structure of the resulting graphs varied dramatically depending on the method used. Importantly, we also show that incomplete genome assemblies (common in large sequencing studies) strongly alter graph structure, and that different tools respond to incomplete data in different ways. In some cases, this affects the detection of medically relevant genes, including Shiga toxin genes linked to severe disease. Our results demonstrate that pangenome graphs are not interchangeable representations of bacterial diversity. Instead, their structure depends on both the method and the quality of the input data. We argue that researchers should choose graph-building tools carefully and report structural properties explicitly to ensure reproducible and interpretable results.

## Introduction

Pangenome graphs provide a flexible framework for representing population-scale genomic diversity by integrating shared and variable sequence into a single structure. In these graphs, homologous sequence elements are represented as nodes, adjacencies and variations across genomes define edges, and individual genomes correspond to paths through the graph[1]. By unifying core and accessory content, pangenome graphs can support tasks that are often treated separately, including gene content analysis and structural variation representation[1–4], and they are increasingly used as datasets scale to thousands of genomes[5,6]. In addition, preserving local genomic context within a unified representation provides a natural substrate for downstream analyses that exploit neighborhood structure, including emerging machine-learning approaches that use graph topology or path structure to model genotype–phenotype relationships[3,7–12]

A wide range of computational strategies has emerged for constructing pangenome graphs, reflecting different assumptions about homology and the resolution at which variation should be represented. Broader overviews of pangenome graph models and construction strategies are available in recent reviews[1,13,14]. In brief, current methods can be grouped into three main paradigms: (1) cluster of orthologous genes (COG) approaches, which build graphs over inferred gene families, producing interpretable gene-centric summaries [3,15–17]; (2) coloured compacted de Bruijn graph (ccDBG) approaches which operate at nucleotide resolution by compacting k-mer graphs into unitigs and annotating nodes and edges with genome membership, capturing fine-scale sequence diversity at population scale [18–23] and (3) multiple sequence alignment (MSA) approaches which derive graphs from multiple sequence alignments or collinear alignment blocks, typically preserving synteny in conserved regions while down-weighting highly divergent segments that are difficult to align reliably [24–27]. These design choices determine which aspects of variation are emphasized or overlooked: COG graphs efficiently summarize coding sequences but typically omit intergenic sequence and can be sensitive to gene-calling differences; ccDBG graphs retain nucleotide-level resolution but can become complex in repeat-rich or paralogous regions; and MSA approaches preserve synteny where alignment is stable but may under-represent highly divergent accessory sequences.

Despite rapid methodological development, the extent to which different tools produce comparable graph representations when applied to the same bacterial genomes remains poorly quantified. Existing benchmarking efforts have already shown substantial variation across pangenome tools in computational performance and in inferred pangenome composition, including differences in the number of genes recovered even when annotation is standardised [28,29], while large-scale evaluations in other domains have highlighted that representation decisions could shape downstream analyses [30,31]. However, these studies have largely evaluated pangenomes in terms of gene content summaries and resource profiles, rather than systematically interrogating how different graph paradigms encode variation at the level of graph topology and connectivity in bacterial datasets. Additionally, although simulation-based studies have explored the effects of fragmentation and incompleteness[3,32], systematic comparisons of bacterial pangenome graph structure using real-world mixtures of draft assemblies and complete genomes remain scarce. This gap is particularly relevant in practice because most microbial genomes are generated and shared as short-read assemblies rather than finished genomes[33], and repetitive and mobile-element–rich regions frequently collapse or fragment under short-read assembly[34–37], altering gene calls and weakening the genomic context signals that many graph methods rely on. As a result, assembly fragmentation may not be a minor nuisance variable but a major determinant of graph topology and pangenome composition, with potential consequences for downstream tasks such as gene presence–absence analyses, population structure inference, and locus-level interpretation in surveillance settings.

Here, we surveyed available approaches for constructing bacterial pangenome graphs and subsequently benchmarked six representative tools spanning COG-, ccDBG-, MSA-based, and hybrid paradigms. Using 175 complete *Escherichia coli* O157:H7 genomes and matched short-read data, we compared graph structure across methods under identical complete-genome inputs and then quantified how graph properties change across controlled mixtures of complete genomes and short-read assemblies. Finally, we assessed recovery of the Shiga toxin loci as a clinically grounded readout of gene-level consequences in repeat- and copy-number–challenging regions. Rather than providing a single overall ranking, our goal is to expose how representation choices and assembly fragmentation jointly shape bacterial pangenome graph properties, and to guide method selection by clarifying the trade-offs, biases, and failure modes that matter for downstream analyses.

## Results

### Differences in graph structures across methods when using complete genomes

We identified six representative pangenome-graph construction tools spanning gene-family–based (Panaroo, PPanGGOLiN), hybrid (ggCaller), compacted de Bruijn graph–based (Bifrost, Cuttlefish2), and MSA-based (Pangraph) approaches (Methods; Supplementary Methods; Supplementary Table S1). All tools were applied to a benchmark dataset of 175 complete *E. coli* O157:H7 genomes (Methods; Supplementary Results S1).

Graph size varied markedly across methods. Node counts ranged from 1,667 (Pangraph) to 117,196 (Cuttlefish2), and edge counts from 2,192 to 157,027, respectively (Table 1).

**Table 1.** Graph features per tool.

| Tool | Method | Node Count | Edge Count | Subgraph Count |
| --- | --- | --- | --- | --- |
| Panaroo | COG | 8740 | 12925 | 6 |
| PPanGGOLiN | COG | 7148 | 9593 | 4 |
| ggCaller | ccDBG+COG | 7625 | 9152 | 582 |
| Bifrost | ccDBG | 58991 | 78439 | 324 |
| Cuttlefish | ccDBG | 117196 | 157027 | 2 |
| Pangraph | MSA | 1667 | 2192 | 332 |

Gene annotation–based methods produced broadly comparable graphs. Panaroo generated slightly more nodes and edges than PPanGGOLiN and ggCaller, but all three showed nearly identical node length distributions (Figure 1A; Supplementary Table S3). PPanGGOLiN did not produce nodes shorter than 100 bp, consistent with its gene-family inference strategy, whereas Panaroo and ggCaller retained shorter elements. Fragmentation differed more substantially: ggCaller yielded 582 subgraphs, with the largest component containing 75.9% of nodes, whereas Panaroo and PPanGGOLiN produced only 6 and 4 subgraphs, respectively, each with >99% of nodes in the largest component (Figure 1B; Supplementary Table S4).

Unitig-based approaches generated the largest graphs. Cuttlefish2 produced more than twice as many nodes and edges as Bifrost, although their node length distributions were nearly indistinguishable (Figure 1A). Maximum node length differed (53,319 bp vs 45,273 bp), and fragmentation patterns contrasted: Bifrost produced 324 subgraphs compared to only 2 for Cuttlefish2. In both cases, however, >98% of nodes resided in the largest component, indicating that fragmentation was largely confined to small peripheral subgraphs.

Pangraph produced the most compact graph and the broadest node length range (16–103,993 bp), reflecting its alignment-based representation of conserved and variable regions. It yielded 322 subgraphs, with the largest component encompassing 73.9% of nodes—the lowest proportion among methods—and a near-linear cumulative subgraph-size distribution (Figure 1B; Supplementary Table S4).

### Differences in how node-degree encodes genomic variation

Node-degree distributions provided insight into how each method encodes genomic variation. Across all graphs, >97% of nodes had degree ≤10, with degree-2 nodes dominating in every tool (Supplementary Figure S2). Bifrost and Cuttlefish exhibited a relative enrichment of degree-3 and degree-4 nodes, whereas Bifrost, ggCaller, and Pangraph showed elevated proportions of isolated (degree-0) nodes, consistent with their higher subgraph counts.

To distinguish how node degree relates to core versus accessory genome representation, we examined degree–prevalence heatmaps normalized within each degree (Figure 1C; Supplementary Table S5). In COG-based graphs (PPanGGOLiN, Panaroo, ggCaller), degree-2 nodes were strongly enriched for high-prevalence segments (≥95% of genomes), indicating that linear paths predominantly represent core genome structure. In PPanGGOLiN and ggCaller, this core bias extended to higher degrees (3–7), suggesting that branch points also frequently encode widely shared sequences. Panaroo showed a more balanced distribution at higher degrees, consistent with greater splitting of gene families according to genomic context. Notably, PPanGGOLiN produced two extremely high-degree nodes (398 and 437), each representing multi-copy gene families present in nearly all genomes (>3,400 total gene instances). BLASTN searches against the NCBI non-redundant database identified these clusters as IS3-family transposases (IS3411 and IS1203).

Unitig-based graphs displayed distinct patterns. In Bifrost, degrees 2–4 encoded rare and core segments in roughly equal proportions, indicating a relatively homogeneous use of node degree. In contrast, Cuttlefish preferentially used degree-2 nodes for low-prevalence accessory segments, while reserving higher degrees (≥3) for core-like unitigs, revealing a clear functional stratification of degree usage.

Pangraph exhibited yet another encoding strategy. Degree-2 nodes were biased towards low-prevalence segments, whereas intermediate degrees (3–4) were more core-associated. Higher degrees (5–7) shifted again toward rare segments, indicating that complex branching structures increasingly represent isolate-specific alignment blocks.

Finally, degree-0 nodes differed markedly across tools. Whereas Panaroo contained only two isolated nodes (both rare), ggCaller, Pangraph, and Bifrost retained substantial numbers of isolated segments, including a non-trivial fraction of high-prevalence sequence. This suggests that, in these graphs, parts of the core genome are not fully integrated into the dominant connected component.

**Figure 1.**
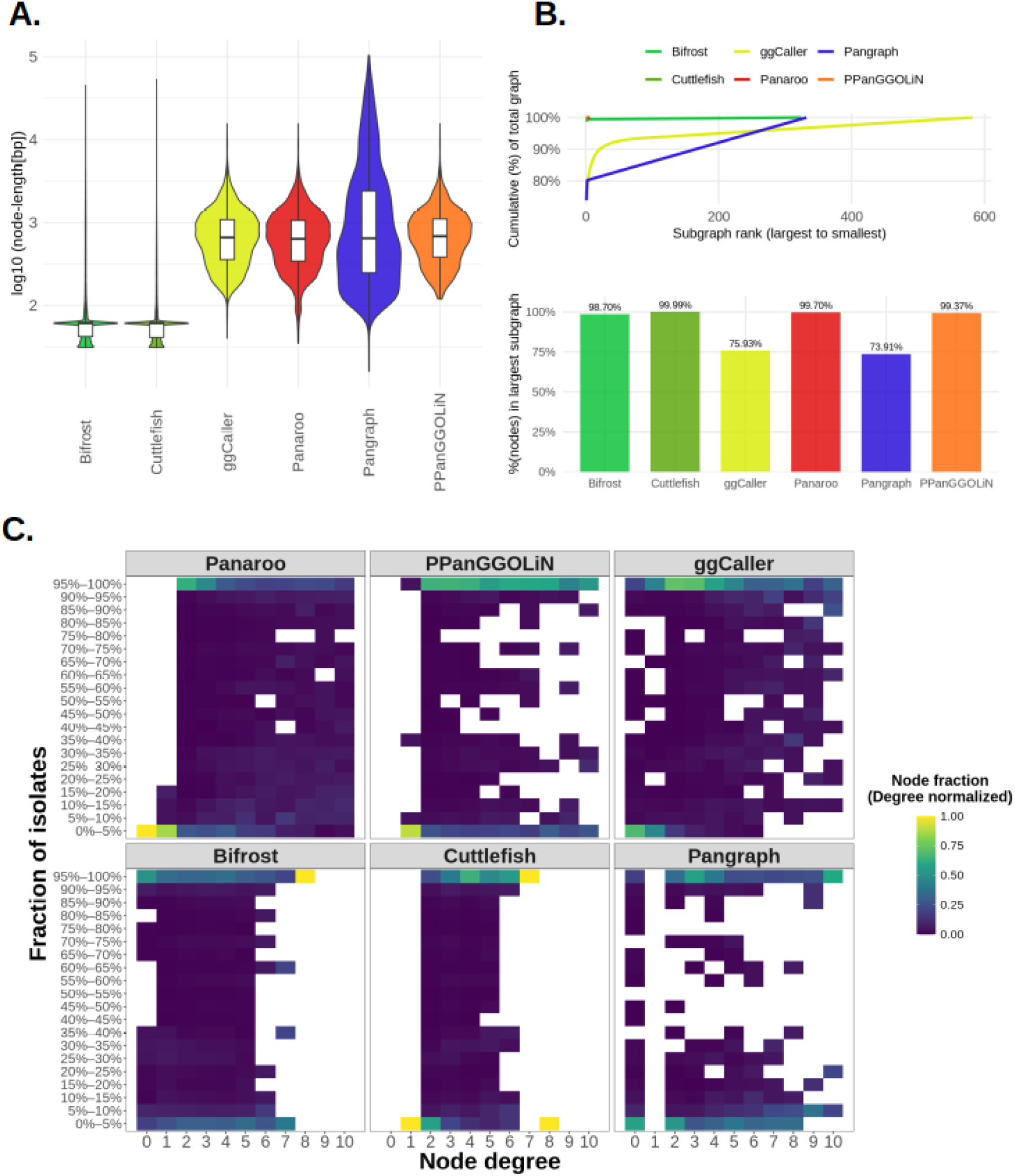
**(A)** Node length distributions for each method, shown as violin plots of log_10_-transformed node length (bp). White boxes indicate the median and interquartile range; whiskers extend to 1.5×IQR. **(B)** Graph fragmentation metrics. Top: cumulative fraction of total graph content captured as subgraphs are added from largest to smallest (subgraph rank). Bottom: fraction of nodes contained in the largest connected component for each method (values shown above bars). **(C)** Degree–prevalence signatures. Heatmaps show node degree (x-axis; degrees 0–10) versus node prevalence across isolates (y-axis; prevalence bins as fraction of isolates). Colours are degree-normalized, i.e., for each tool and degree, the colour represents the fraction of nodes of that degree falling into each prevalence bin (rows within a degree sum to 1).

### Impacts of genome fragmentation

#### a) Global graph structure and connectivity

To evaluate the effect of genome fragmentation, we reconstructed pangenome graphs from mixtures of complete and draft assemblies and expressed all metrics as percentage change relative to the 100% complete baseline (Figure 2; Supplementary Table S6). Pangraph was excluded because it did not run reliably on fragmented datasets.

Fragmentation produced paradigm-dependent effects on graph size. Among COG-based tools, increasing assembly fragmentation generally reduced graph complexity. Panaroo showed the strongest contraction, with edge counts decreasing by ∼40% and node counts by ∼15–20% in the fully draft dataset. PPanGGOLiN was comparatively stable, with minimal changes in node counts and modest reductions in edges (≤10%). ggCaller exhibited intermediate behavior, with moderate edge reductions (∼15%) and only minor variation in node counts.

In contrast, unitig-based tools displayed inflation under fragmentation. Bifrost showed modest increases (<10%) in node and edge counts, whereas Cuttlefish exhibited substantially stronger inflation (≈20–30% in the fully draft dataset), which decreased progressively as the proportion of complete genomes increased.

Fragmentation also markedly affected graph connectivity. Subgraph counts increased progressively as the proportion of draft assemblies rose (Figure 2). Panaroo and PPanGGOLiN were particularly sensitive, with subgraphs rising from 6 to 145 (≈+2300%) and from 4 to 66 (≈+1550%), respectively, in the fully fragmented dataset. Bifrost and ggCaller showed more moderate increases, from 324 to 498 (≈+54%) and from 582 to 723 (≈+24%), respectively. Cuttlefish, despite starting from only two subgraphs, increased to six (≈+200%). Across tools, fragmentation consistently promoted graph disconnection, although the magnitude of this effect varied substantially by method.

#### b) Node-degree usage and core vs accessory representation

We examined how fragmentation redistributes graph mass across node degrees by contrasting fully complete and fully draft datasets (Supplementary Figure S3). In COG-based graphs, fragmentation generally shifted mass from higher-degree nodes toward lower-degree ones. In Panaroo and ggCaller, degree-0 and degree-1 nodes increased and degree-2 showed the largest enrichment, whereas degrees ≥3 were consistently depleted.PPanGGOLiN showed the same overall shift toward lower degrees, but with enrichment concentrated in degree-1 and a slight decrease in degree-2. Together, these patterns indicate that fragmented input simplifies complex junctions into linear or terminal segments.

**Figure 2.**
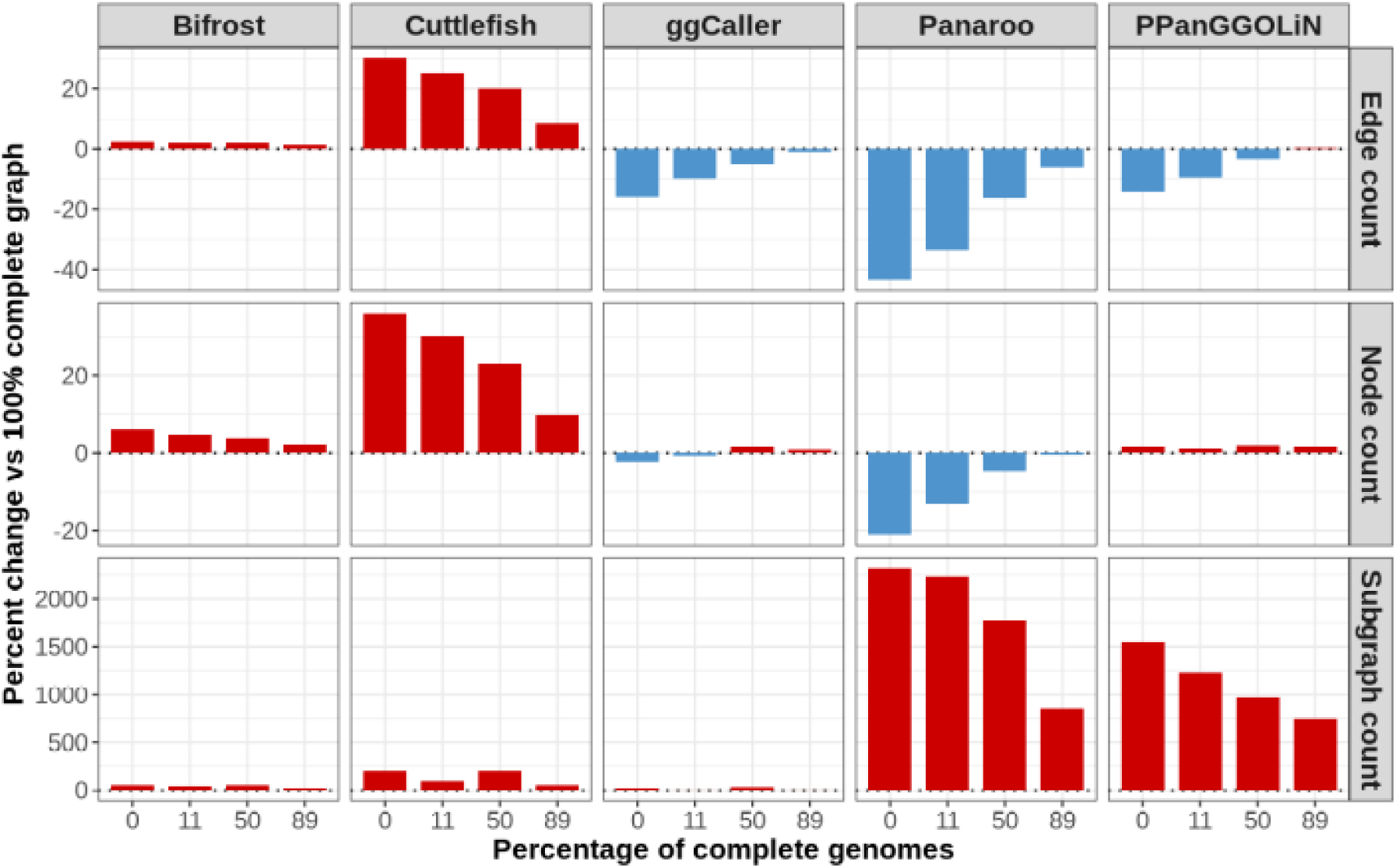
Pangenome graphs were reconstructed from mixtures of complete genomes and fragmented short-read assemblies, and graph metrics are reported as percent change relative to the graph built from 100% complete genomes. Pangraph was excluded because it did not run reliably on fragmented-assembly datasets. Facet columns indicate graph-construction methods and x-axes denote the percentage of complete genomes in each mixture. Facet rows show percent changes in edge count (top), node count (middle), and subgraph count (bottom). Positive values (red) indicate inflation relative to the 100% complete genomes reference graph (dotted line), whereas negative values (blue) indicate reductions.

Unitig-based methods showed a related but distinct redistribution. In both Bifrost and Cuttlefish, degree-1 nodes increased markedly under fragmentation. In Bifrost, this was accompanied by a broad depletion across degrees ≥2, indicating a general simplification of branching structure. In contrast, Cuttlefish exhibited a more structured shift: degree-2 and degree-4 nodes were depleted, whereas degree-3 nodes increased, suggesting that fragmentation is accommodated not only by generating terminal unitigs but also by introducing additional low-order branch points.

Because the relative size of the core and accessory genome underpins biological interpretation and downstream comparative analyses, we next examined how fragmentation alters this balance (Figure 3, Supplementary table S7). For most tools, fragmentation reduced the fraction of nodes classified as core (95–100% prevalence) and increased the accessory fraction (0–5%).

PPanGGOLiN and ggCaller showed comparable shifts, with core depletion concentrated at degrees 2–4 and accessory enrichment primarily in degree-0 and degree-1 nodes. Thus, under fragmented input, accessory regions are increasingly encoded as isolated nodes and terminal segments, while shared junction structures that normally represent core genome continuity are diminished. Panaroo exhibited the opposite trend: the fraction of nodes classified as core increased under fragmentation, largely through enrichment of degree-2 nodes in the core bin. Concurrently, accessory degree-2 nodes were depleted. This suggests that Panaroo reinforces a simplified degree-2 backbone when confronted with fragmented assemblies.

Unitig-based methods mirrored several of these trends but with distinct signatures. In both Bifrost and Cuttlefish, the global fraction of core nodes declined while accessory fractions increased under fragmentation. In both tools, core depletion was concentrated at degrees 2–4, indicating that shared backbone continuity is reduced. In Bifrost, this loss was partially offset by an increase in degree-1 core unitigs, suggesting that some core segments are re-encoded as terminal nodes. The representation of the accessory genome diverged between the two tools. In Bifrost, accessory gains were distributed across degrees 1–4. In contrast, Cuttlefish showed accessory enrichment primarily in degree-1 and degree-3 nodes, coupled with depletion of degree-2 accessory unitigs.

#### c) Computational cost

We quantified the effect of fragmentation on computational performance by recording total CPU time and peak resident memory for each tool on the 0%, 50%, and 100% complete-genome datasets (Figure 4).

Among COG-based methods, Panaroo and PPanGGOLiN showed comparable CPU requirements (≈3–6×10³ CPU-seconds), with higher proportions of complete genomes associated with modest increases in runtime (Figure 4A). ggCaller was consistently more computationally demanding, increasing from 1.9×10⁴ CPU-seconds in the all-draft dataset to 7.6×10⁴ CPU-seconds in the fully complete dataset.

Unitig-based tools displayed contrasting behavior. Bifrost was the fastest overall (≈5–7×10² CPU-seconds) and showed minimal sensitivity to fragmentation. In contrast, Cuttlefish exhibited extreme fragmentation dependence: CPU time ranged from 1.3×10⁴ CPU-seconds in the fully complete dataset to 2.1×10⁶ CPU-seconds in the all-draft condition, nearly a three-order-of-magnitude increase.

Peak memory usage varied less than CPU time (Figure 4B). Panaroo required 3.1–6.0 MB of RAM, PPanGGOLiN 7.3–7.7 MB, and ggCaller 29–33 MB. Bifrost and Cuttlefish used <0.2 MB across all datasets.

**Figure 3.**
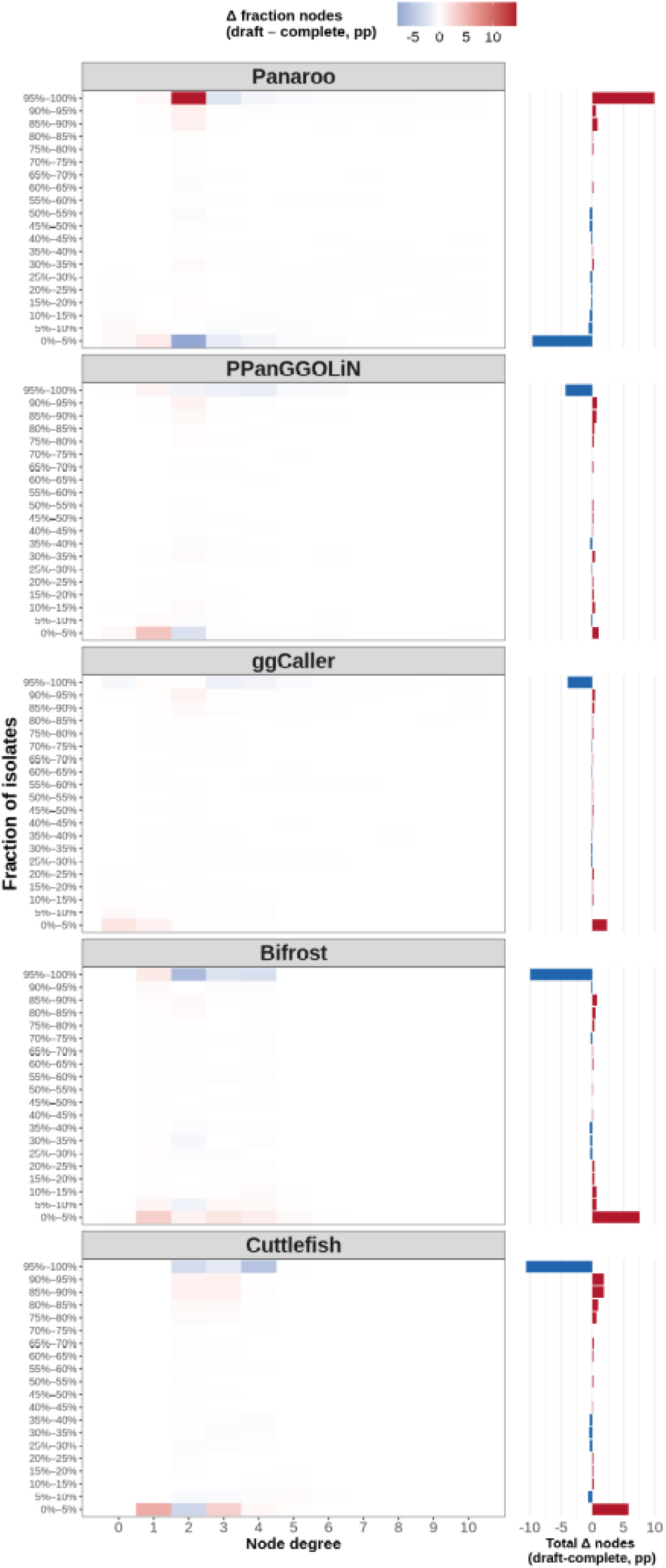
Heatmaps show how fragmentation (fully draft vs fully complete genomes) redistributes nodes across degree (0–10; x-axis) and prevalence bins (fraction of isolates; y-axis). Colours represent the percentage-point difference in node fraction (draft − complete): red indicates enrichment in the draft graph and blue depletion. Fractions are normalized to the total number of nodes per graph. The bar plot (right) summarizes total changes across prevalence bins.

**Figure 4:**
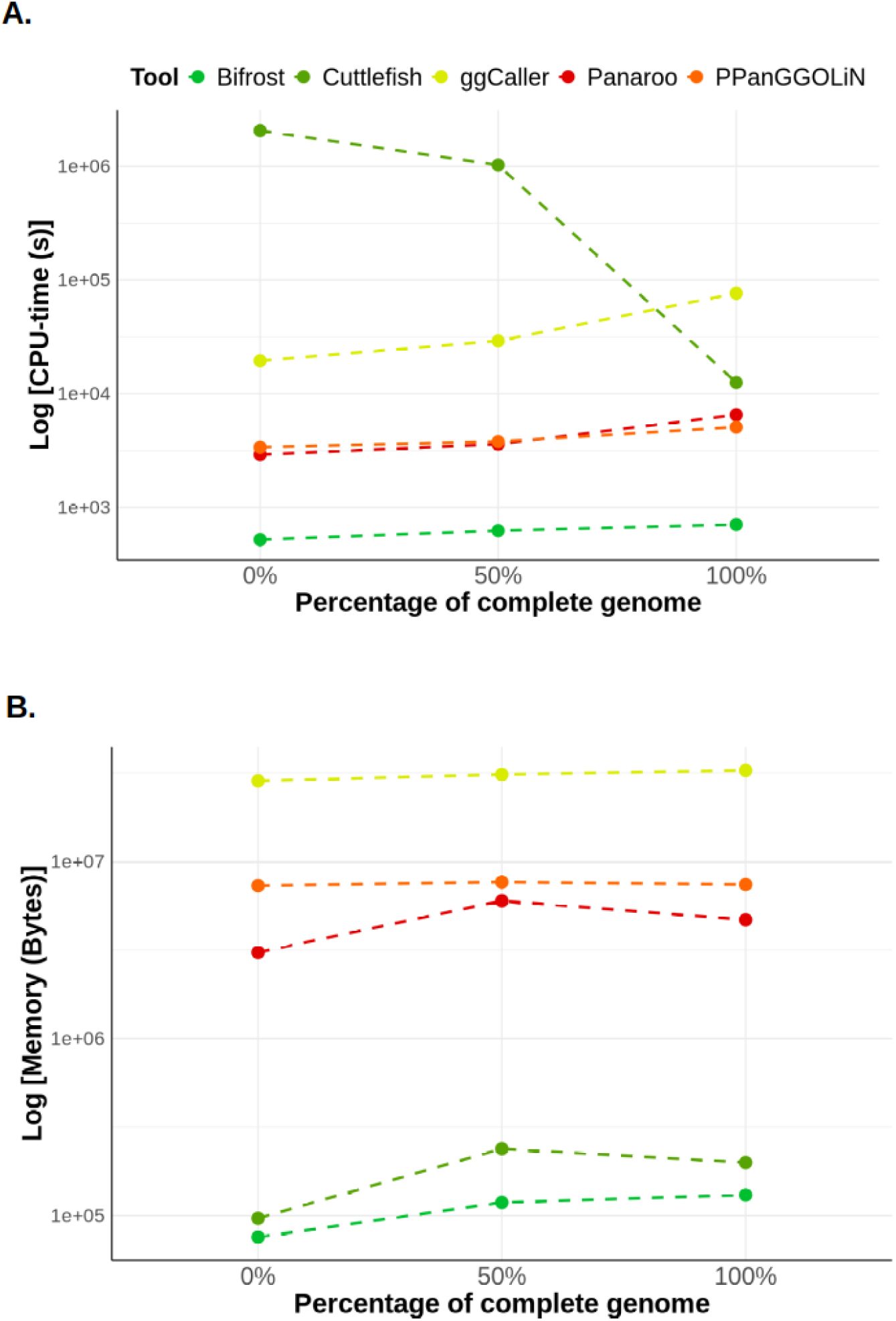
Total CPU time **(A)** and peak memory usage **(B)** by the evaluated tools as a function of the percentage of complete genomes included in pangenome graph construction.

### Calling *stx* genes

Because short-read assemblies frequently fragment or collapse the *stx* locus, particularly in isolates carrying multiple toxin variants, we used *stx* as a focused benchmark to evaluate each method’s ability to detect and type clinically relevant genes (Supplementary Figure S4; Supplementary Table S8).

This analysis was restricted to COG-based methods that report gene calls directly. Performance was assessed using recall and precision (Figure 5), stratified by assembly status (Complete, Fragmented, Collapsed) and toxin subunit (*stxA* and *stxB*). Bakta annotations were included as a baseline, representing per-isolate gene calling without pangenome-level reconciliation.

Across the dataset, the ground truth comprised 266 *stx* operons. Most copies were Complete, with substantial fractions classified as Fragmented and a smaller subset as Collapsed. Fragmentation and collapse were enriched in multi-variant or multi-copy profiles (Supplementary Results S2), indicating that assembly artefacts concentrate in precisely those backgrounds where accurate typing is most challenging.

For *stxA*, Panaroo and PPanGGOLiN maintained perfect precision across all strata but did not improve recall relative to Bakta (Figure 5A). Recall remained constrained in Fragmented and Collapsed loci even when using 100% complete genomes. In contrast, ggCaller recovered near-complete recall in Complete and Collapsed strata once ≥50% complete genomes were included, and matched Bakta for fragmented loci at high completeness. However, this improvement came at the cost of reduced precision, particularly for Complete and Fragmented loci.

Results for *stxB* were broadly similar, with performance strongly dependent on assembly status (Figure 5B). Precision remained near-maximal across tools, with only minor decreases for ggCaller in Complete loci. Under fragmentation, ggCaller and Panaroo achieved recall comparable to Bakta across mixtures, whereas PPanGGOLiN showed persistently lower recall. For Collapsed loci, all pangenome methods remained below the Bakta baseline, indicating limited recovery of missing copies in collapse-prone backgrounds (Supplementary Results S2).

**Figure 5:**
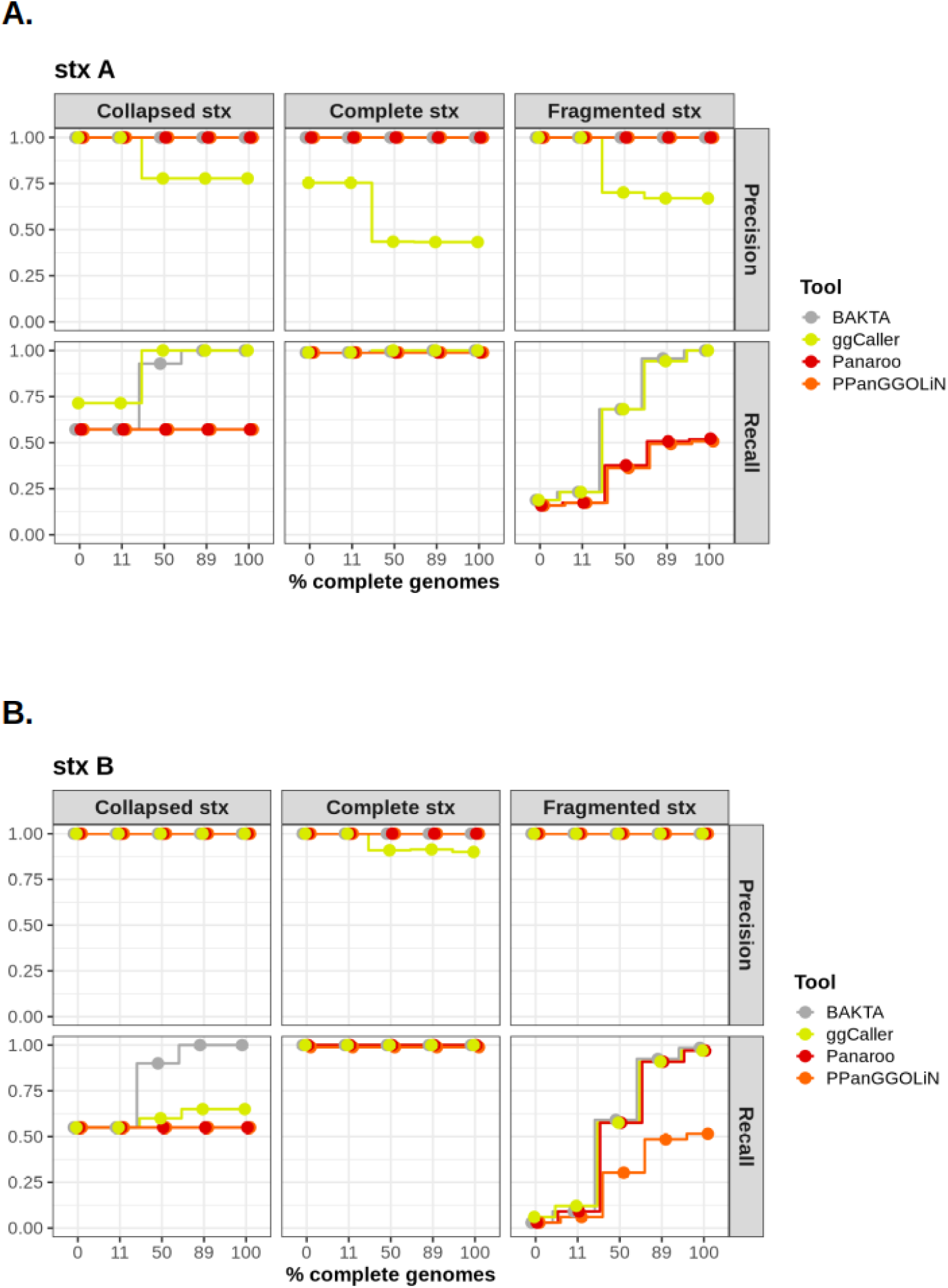
Precision and recall are shown for the detection of the shiga-toxin operon subunit-A **(A)** and subunit-B **(B)** as the fraction of complete genomes in the dataset increases (x-axis). Performance is stratified by *stx* locus assembly status in the short-read assemblies (Collapsed, Complete, Fragmented). Curves compare COG-based pangenome methods (Panaroo, PPanGGOLiN, ggCaller) against BAKTA as a baseline per-isolate annotation approach. Metrics were computed against the ground-truth copy set derived from calling stx operons using VirulenceFinder in complete genomes.

## Discussion

In this work we identified pangenome-graph building methods spanning COG-, ccDBG-, MSA-based, and hybrid paradigms, and compared representative tools across these classes. Differences across tools should not be interpreted as implementation detail alone; they reflect distinct operational definitions of homology, variation, and genomic context that constrain downstream inference. Recent benchmarking in clonal bacteria supports this framing: even when true gene-content diversity is limited, pan-genome size estimates can vary substantially across COG pipelines due to annotation behaviour and assembly characteristics[38].

We designed the evaluation around a stringent and practically relevant setting: a repeat-rich, clinically important STEC lineage where short-read assemblies are expected to fail in precisely the regions that matter most for assessing pathogenic potential and clinically relevant determinants[37,39–43]. Our O157:H7 dataset is dominated by ST11, reducing phylogenetic heterogeneity while preserving substantial within-lineage accessory genome variation. Elevated repeat content, frequent multi-variant toxin carriage and the concentration of key determinants in mobile, repeat-dense compartments (including prophages and other accessory elements), create a scenario in which graph methods must contend with repeats and mosaicism while still supporting locus-level ground truth for clinically relevant genes.

When applied to identical complete genomes, methods differed by orders of magnitude in size and fragmentation, indicating that global topology is primarily determined by representation strategy. ccDBG-based methods produced the largest graphs (up to 117,196 nodes with Cuttlefish), consistent with nucleotide-resolution graphs retaining local variants typically collapsed by gene-family approaches. Pangraph produced the most compact graph (1,667 nodes) but with comparatively strong fragmentation, suggesting that MSA-based approaches can aggressively collapse homologous regions yet may fail to connect highly divergent accessory sequences into a single coherent structure. This behaviour is plausibly tunable via Pangraph’s similarity/merging thresholds, but exploring that parameter space was beyond the scope of this benchmark.

Within ccDBG graphs, Bifrost and Cuttlefish yielded nearly identical node-length distributions yet diverged markedly in node and edge count, with Cuttlefish producing roughly twice as many nodes and edges as Bifrost. This is consistent with distinct compaction formalisms over the same k-mer space: Cuttlefish’s state-based formulation determines unitig boundaries via global side-classification and yields many explicit unitigs/adjacencies[19], whereas Bifrost’s construction and correction pipeline favours continuity during compaction which could yield fewer nodes/edges[18]. Notably, higher continuity does not necessarily imply higher global connectivity: Cuttlefish produced no degree-0 nodes in our exports, whereas Bifrost generated many degree-0 unitigs that were nevertheless highly prevalent (95–100% of genomes), inflating the number of connected components without substantially affecting the main backbone. A parsimonious interpretation is that Bifrost can retain ubiquitous sequences as intact unitigs while materialising fewer junction adjacencies when flanking context is ambiguous across genomes, leaving some shared sequence topologically isolated. Further work is needed to determine the biological interpretation of these isolated core-unitigs.

By contrast, COG-based approaches generated graphs of broadly comparable size, with connectivity representing the most salient divergence. ggCaller generated hundreds of subgraphs, whereas Panaroo and PPanGGOLiN produced graphs in which >99% of nodes fell within the largest component. Given ggCaller’s workflow (gene families inferred on top of a Bifrost-derived ccDBG substrate), fragmentation in the upstream graph can propagate into the final COG-based graph. Beyond connectivity, differences in node and edge counts among COG tools reflect distinct heuristics for collapsing versus splitting orthologous groups[3,16], including paralog handling and the degree to which neighbourhood/context is used during correction. This is consistent with the observed degree–prevalence signatures: PPanGGOLiN and ggCaller show strong enrichment of core nodes across degrees 2–7, whereas Panaroo exhibits a more balanced rare-versus-core distribution at higher degrees (notably degrees 3–4), consistent with stronger context-driven splitting rather than collapsing into multi-genome “core” junction structures.

A central result is that assembly fragmentation is a first-order determinant of graph structure, with effects that are directionally consistent within paradigms. In COG-based graphs, replacing complete genomes with drafts reduced graph size, most clearly through edge loss. This contraction was strongest in Panaroo (≈40% fewer edges), modest in PPanGGOLiN (≈5–10%), and intermediate in ggCaller (≈15%). Draft assemblies collapse repeats, truncate genes, and disrupt neighbourhood context, reducing both effective gene recovery and adjacency evidence[32,44,45]. Annotation discrepancies can amplify these effects, inflating apparent accessory variation and depressing core recovery[38]. Panaroo’s larger contraction aligns with its design goal of pruning contig-end artefacts[3], whereas PPanGGOLiN’s relative stability reflects inference driven primarily by cross-genome presence/absence patterns, with neighbourhood used as smoothing rather than a prerequisite for family definition[16]. Unitig graphs expanded under fragmentation, consistent with contig breaks introducing additional unitig boundaries and reshaping junction structure. Bifrost showed modest growth (<10%), whereas Cuttlefish expanded more strongly (∼20–30%), indicating greater sensitivity of its compaction strategy to continuity loss. Degree–prevalence profiles suggest that fragmentation increases terminal structure in Bifrost, while more substantially reshaping junction structure in Cuttlefish. Thus, draft-input topology is not merely a scaled version of the complete-genome graph; contig breaks propagate differently into unitig boundaries and junction order across implementations, with potential downstream consequences.

Computational trends broadly mirrored structural effects. For all methods except Cuttlefish, CPU time increased with the proportion of complete genomes, consistent with larger and more connected graphs requiring greater reconciliation effort[32]. Cuttlefish was the exception, with CPU time peaking on all-draft inputs and dropping sharply for all-complete, consistent with draft assemblies inducing larger unitig graphs, with a higher number of unitigs/edges and costly boundary/state decisions[19]. These comparisons reflect graph construction given fixed inputs; COG-based timings exclude upstream annotation, and our intent was not to rank pipelines but to quantify how fragmentation modulates runtime within each representation class.

The stx locus analysis provides a clinically meaningful readout of how representation and completeness effects translate into gene-level outcomes. Panaroo and PPanGGOLiN maintained perfect precision for *stxA* but did not improve recall beyond the Bakta baseline, with recall remaining constrained in fragmented and collapsed contexts. ggCaller uniquely recovered near-complete recall in collapsed and complete strata once ≥50% complete genomes were present, but at the cost of reduced precision. This pattern is consistent with ggCaller’s design goal of mitigating contig-break artefacts by performing gene calling directly on a population de Bruijn graph: sequence paths present in other genomes can bridge local breaks in a given assembly, increasing the chance of recovering full-length loci that would otherwise be split or partially called[32]. At the same time, graph-based recovery can increase ambiguity around gene boundaries (particularly for short open reading frames or repeat-associated regions), providing a plausible route to inflated false positives and the precision drop we observed[32]. For *stxB*, precision was generally near maximal, but recall remained sensitive to assembly status and method, including persistent under-recovery of collapsed copies. These results show that pangenome-level reconciliation does not automatically correct assembly artefacts at clinically relevant loci; instead, tools occupy distinct precision–recall trade-offs that depend on locus status and dataset completeness. The divergence between *stxA* and *stxB* further indicates locus dependence even within the same operon, and other genes may exhibit distinct failure modes, consistent with prior observations that fragmentation and contig-end effects can generate partial calls and missed genes in pangenome workflows[3,32]. Finally, while nucleotide-graph and MSA-based approaches should in principle support recovery of *stx* variants by coupling their output to sequence-to-graph aligners[46–48], doing so would add an additional mapping layer whose sensitivity and bias can vary; benchmarking that end-to-end stack was beyond the scope of this study.

Several limitations bound inference while pointing to next steps. The dataset is dominated by ST11 O157:H7, chosen to isolate methodological effects under a repeat-rich background; results may differ in species with lower repeat load or alternative recombination dynamics[3]. Our locus-level evaluation excludes unitig graphs without an additional mapping layer, and Pangraph could not be compared across fragmentation gradients due to failures on draft assemblies. Future benchmarks should standardise gene-translation steps for nucleotide graphs, expand to multi-species panels, and evaluate downstream tasks under explicit completeness gradients. Annotation policy remains an additional axis: differences in CDS calling can systematically shift inferred core/accessory composition even when nucleotide sequence is conserved[38]. Prior benchmarking in bacteria has likewise shown that inferred pangenome composition can be sensitive to tool choice[28,29], and benchmarks in other domains emphasise evaluating representations in the context of intended downstream tasks rather than solely by size or runtime[30,31].

Two practical conclusions follow. First, method choice must be task-driven: gene-cluster graphs provide interpretable summaries but exhibit completeness-dependent connectivity changes; unitig graphs preserve nucleotide diversity but inflate under fragmentation and may incur computational penalties; and MSA approaches may offer compact representations yet struggle operationally on fragmented datasets. Second, completeness composition is a primary design variable: mixing complete and draft genomes is not neutral, and the same cohort can yield systematically different graphs as complete/draft genome mixture changes.

In summary, bacterial pangenome graphs are representation-dependent models rather than interchangeable outputs. Assembly fragmentation is a primary driver of topology, and different paradigms respond to incompleteness in systematically distinct ways. Method choice must therefore be task-driven and completeness-aware. Making graph structure, scalability, and locus-level failure modes explicit is essential for reproducible downstream inference and correct interpretation of bacterial pangenomes.

## Methods

### Survey of available methods to construct pangenome graphs

Available pangenome construction methods were surveyed by conducting a search in the PubMed database using the following terms: ((((((((((((((pangenome[Title/Abstract]) OR (pan-genome[Title/Abstract])) OR (pangenomics[Title/Abstract])) OR (pan-genomes[Title/Abstract])) OR (pan-genomics[Title/Abstract])) OR (pangenomes[Title/Abstract])) OR (core-genome[Title/Abstract])) OR (core genome[Title/Abstract])) OR (accessory genome[Title/Abstract])) OR (accessory-genome[Title/Abstract])) OR (compacted de Bruijn[Title/Abstract])) OR (coloured compacted de Bruijn[Title/Abstract])) OR (colored compacted de Bruijn[Title/Abstract])) AND ((((((graphical[Title/Abstract]) OR (graph[Title/Abstract])) OR (network[Title/Abstract])) OR (graph-based[Title/Abstract])) OR (graphs[Title/Abstract])) OR (graphical structure[Title/Abstract]))) AND ((((((((((((pipeline[Title/Abstract]) OR (tool[Title/Abstract])) OR (package[Title/Abstract])) OR (method[Title/Abstract])) OR (software[Title/Abstract])) OR (library[Title/Abstract])) OR (model[Title/Abstract])) OR (toolkit[Title/Abstract])) OR (algorithm[Title/Abstract])) OR (algorithms[Title/Abstract])) OR (algorithmic[Title/Abstract])) OR (data structure[Title/Abstract]))

On November 1^st^ 2024, this search criteria returned 223 publications, which were manually curated to find a total of 14 relevant pangenome-graph construction tools, which were grouped according to their underlying graph-construction strategy: three relied on clustering homologous genes into COGs, five used ccDBGs, four employed MSA, and two combined COG identification with ccDBG construction (Supplementary Table S1). From these, we selected six tools for downstream comparison: Panaroo (COG), PPanGGOLiN (COG), ggCaller (ccDBG + COG), Bifrost (ccDBG), Cuttlefish2 (ccDBG), and Pangraph (MSA). A detailed description on rationale behind tool selection for benchmarking efforts can be found in Supplementary Methods. A total of 28 tools were excluded from downstream analyses due to diverse reasons, which are listed in Supplementary Table S9.

### Retrieving *E. coli* O157:H7 complete genomes and short reads from public databases

Ncbi-genome-download v0.2.10 was used to download all *E. coli* sequences labeled as ‘complete genomes’ up to 30th October 2023 (n = 2980) (https://github.com/kblin/ncbi-genome-download/). *In silico* serotyping was performed using ECTyper[49] v1.0. Sequence types (STs) were determined using mlst v2.23.0 (https://github.com/tseemann/mlst). Virulence-gene content was determined using Virulence Finder[50] v2.0.4.

The metadata of the isolates was retrieved and parsed using Entrez-utilities[51] v13.9. BioSample IDs were used as input for downloading Illumina raw reads corresponding to the complete genomes by using SRA Tools v3.0.9)(https://github.com/ncbi/sra-tools) .

### Quality control of short-read data and genome assembly

Adapter contamination and bases with a phred-score below 20 were removed using Trim Galore v0.6.10 (https://github.com/FelixKrueger/TrimGalore). Fastqc v0.12.1 was used for quality control of pre- and post-trimming samples. Unicycler[52] v0.5.0 with default parameters was used for genome assembly. Short-read assembly quality was assessed by aligning these against the corresponding complete genome using QUAST[53] v5.2.0. Genomes with a fraction of misassembled and unaligned contigs greater than 4.0% were removed (n=4). CheckM2[54] v1.10 was applied to evaluate the quality of remaining genomes (n=175), with all displaying completeness above 99.0% and contamination smaller than 1.0% .

### Benchmark dataset analysis

A neighbor-joining phylogeny was generated to contextualize the diversity of the *E. coli* isolates included in the analysis. All complete *E. coli* genomes available in RefSeq at the time of study were processed using Mashtree[55] v1.4.6, which computes pairwise Mash[56] distances and constructs a neighbor-joining tree from the resulting distance matrix. Default parameters were used. The resulting phylogeny was uploaded to Microreact[57] for visualization and integration with serotype, *stx* profiles and plasmid metadata. An interactive version of the tree is publicly available at: https://microreact.org/project/stec-benchmark.

Plasmids from the benchmark dataset were assigned to clusters using mge-cluster[58] v1.1 coupled to the existing *E. coli* plasmid model, which can be accessed at: https://doi.org/10.6084/m9.figshare.21674078.v1.

K-mer counts for all isolates were obtained using Jellyfish[59] v2.3.1 . For each genome, canonical k-mers were counted using the jellyfish count command (k = 21).

Summary statistics were extracted with the jellyfish stats function, which reports (i) the number of unique k-mers (k-mers occurring exactly once), (ii) the number of distinct k-mers (the cardinality of the k-mer set, irrespective of multiplicity), and (iii) the total number of k-mers observed, including all repeated instances. These definitions follow the Jellyfish documentation, available at: https://github.com/gmarcais/Jellyfish/blob/master/doc/Readme.md

To quantify the repeat burden of each genome, we calculated the fraction of repeated k-mer instances, defined as the proportion of all k-mer occurrences in the genome that originate from repeated regions. For a given genome i this metric is computed as detailed in equation 1.

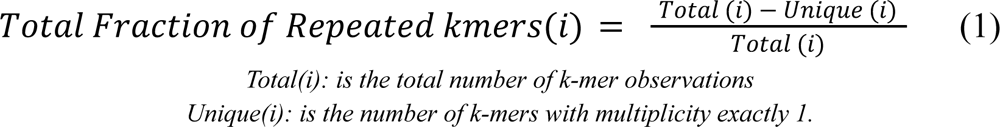

Because the denominator in equation 1 corresponds to all k-mer instances originating from sequences that occur more than once in the genome, the metric *“Total fraction of repeated k-mers”* represents the proportion of the genome composed of repeated elements at the k-mer scale. Higher values indicate genomes with a greater abundance of repeated sequences. This metric provides a genome-wide estimate of repetitiveness informative for evaluating pangenome graph construction, as repeated elements often contribute to graph fragmentation and structural complexity.

### Selecting samples to create mix-datasets, containing both complete and draft-genomes

Five datasets, each containing 175 genomes, were constructed to evaluate the impact of genome fragmentation on pangenome graph structure. These datasets differed only in the proportion of complete genomes versus draft assemblies: 0%, 11%, 50%, 89%, and 100% (corresponding to 0, 20, 88, 156, and 175 complete genomes, respectively; counts rounded to the nearest integer). To choose which genomes were complete in each mixed dataset, we followed what is described here[44]. Briefly, Jaccard distances were computed from a Panaroo gene presence–absence matrix derived from draft assemblies. We embedded the distances with t-SNE and applied k-means clustering to the 2-D embedding, setting “k” equal to the intended number of complete genomes for that dataset. For each cluster, the genome closest to the centroid was designated as a complete genome; the remaining genomes were included as draft assemblies. The proportions of complete genomes selected using this method were 11%, in the rest of the cases, complete genomes were selected randomly. Using 11% as a fraction of complete genomes was motivated by the maximum number of clusters identified by this procedure (n=20) (Supplementary Figure S4). A full list of all complete- and draft-genome combinations can be found in Supplementary Table S10.

### Pangenome graph construction and data parsing

All pangenome graph construction tools were executed with default parameters on a high-performance computing cluster. For tools that require genome annotation files as input, the annotation files were created using Bakta[60] v1.9.1. Parsing and analysis of pangenome graphs were performed using the igraph[61] v0.10.13 and NetworkX[62] v3.4 packages in Python v3.9.18, using in-house scripts, available in the git repository associated to this work.

### Degree-wise analyses of node abundance and prevalence

We performed three complementary degree-wise analyses to quantify how different tools encode genomic variation and how genome fragmentation affects graph structure. First, we compared node degree distributions between graphs. Second, we examined degree-normalised prevalence profiles to assess how each degree is used to represent core versus accessory regions. Finally, we used whole-graph-normalised prevalence profiles to study how core- and accessory-genome representations change when comparing graphs built from complete genomes and from draft assemblies. For each tool *t*, dataset type *d* (Complete or Draft), and node degree *k*, we counted the number of nodes *N_t,d,k_* with degree *k* and computed the total number of nodes in that graph, as detailed in equation 2:

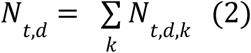

We then expressed degree-wise abundance as the fraction of nodes of that degree within the graph according to equation 3:

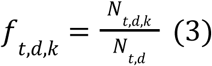

so that 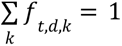 for each (t,d). These fractions were used to describe how each tool distributes graph mass across degrees, as represented in Supplementary Figure S1. To quantify how genome fragmentation redistributes graph mass across node degrees, we calculated, for each tool and degree, the change in node fraction between the draft-only and complete-only graphs, expressed in percentage points, as detailed in equation 4.

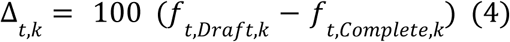

Positive values of Δ*_t,k_* indicate that a larger fraction of the graph’s nodes have degree *k* in the draft-based graph than in the complete-based graph (i.e. fragmentation enriches that degree), whereas negative values indicate depletion, as indicated in Supplementary Figure S3.

To characterise how different degrees encode core versus accessory regions, we further stratified nodes by their prevalence across isolates. For each tool *t*, dataset *d,* degree *k* and prevalence bin *b* (5% intervals from 0–5% to 95–100%), we counted the number of nodes N_t,d,k,b_ whose prevalence fell within bin b and computed the total number of nodes of that degree (equation 5):

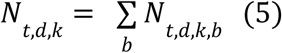

We then derived a degree-normalised prevalence profile (equation 6):

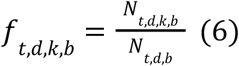

so that 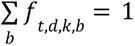 for each (t,d,k). These degree-normalised profiles describe, for a given tool and degree, how nodes are distributed along the core–accessory spectrum, as displayed in Figure 2C.

To assess how fragmentation alters the *overall* use of each degree along this spectrum, we additionally computed a whole-graph-normalised prevalence profile. For each tool t and dataset d, we calculated the total number of nodes in the graph (equation 7)

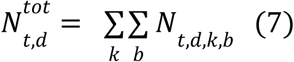

and defined the corresponding fraction (equation 8)

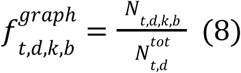

such that 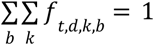. The impact of fragmentation was then summarised as a percentage-point difference between draft and complete graphs (equation 9):

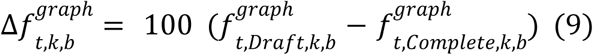

Positive values of Δf_t,k,b_ indicate that a larger proportion of nodes fall into that particular prevalence bin (b) and node-degree (k) combination in the draft-based graph than in the complete-based graph, whereas negative values indicate depletion, as displayed in Figure 4.

### Runtime and memory profiling

To quantify the computational impact of genome fragmentation, we measured runtime and peak memory usage for each tool and completeness setting (0%, 50%, and 100% complete genomes) on the same HPC cluster used for the graph analyses. All jobs were submitted through the SLURM workload manager with identical resource requests per tool. For each run, CPU time (in seconds) and maximum resident set size were obtained from SLURM accounting records (CPUTime and MaxRSS fields, via sacct command). Peak RAM usage reported in the Results corresponds to this MaxRSS value, converted to bytes, while runtime corresponds to total CPU time. For visualisation and comparison across tools, both CPU time and peak RAM were transformed using log_10_ scales.

### *Stx*-content determination

The reference (*ground-truth*) *stx* genotype of each isolate was defined from the corresponding complete genome using VirulenceFinder[50] v2.0.4. These VirulenceFinder calls were used as the gold standard for evaluating *stx* detection and typing in short-read–derived pangenome graphs.

To characterise assembly artefacts at the *stx* locus in short-read assemblies, we identified *stx* gene hits and partial matches using BLASTN[63] v2.15.0. A custom BLAST database was built from the *stx* allele sequences referenced by VirulenceFinder. For each expected *stx* operon copy (A and B subunits), BLASTN alignments (99% identity, 10% coverage) were used to classify the short-read locus as Complete, Fragmented, or Collapsed. A locus was considered “complete” when a single contiguous hit covered the expected allele; “fragmented” when the allele was split across multiple contigs/partial hits; and “collapsed” when multiple expected copies/alleles were represented by a single assembled locus consistent with copy-number collapse (criteria described in Supplementary Table S8).

For each pangenome method (Panaroo, ggCaller, PPanGGOLiN), *stx* gene calls were extracted from the resulting gene clusters using in-house Python scripts. Because *stx* operons are encoded by two subunits (A and B), performance was evaluated separately for stx_A and stx_B, which typically reside in distinct gene clusters. For each isolate, predicted *stx* subunit presence/absence (and variant where applicable) was compared to the gold standard to count true positives (a gold-standard *stx* subunit correctly recovered), false positives (a predicted *stx* subunit absent from the gold standard), and false negatives (a gold-standard *stx* subunit not recovered). These counts were then used to compute precision and recall as defined in Equations 10A and 10B.

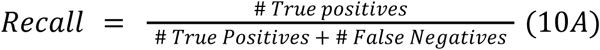

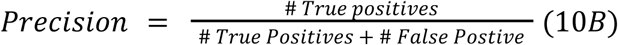

## Data availability Statement

All codes necessary to reproduce the result of this work can be found in the following git repository: https://github.com/jpaganini/benchmarking_pangenome_tools

If required, all raw pangenome graphs can be downloaded from: https://doi.org/10.5281/zenodo.18405233

Phylogenetic tree displayed in Supplementary Figure S1 can be accessed at: https://microreact.org/project/stec-benchmark

Supplementary Materials and Tables can be found in the online version of this article.

## Author Contributions

Conceptualization: JAP; Data Curation: PL and JAP; Formal Analysis: PL, KH, JAP; Funding Acquisition: TJD; Methodology: PL, KH, TJD and JAP; Project Administration: JAP, LM-G, TJD; Software: PL and JAP; Supervision: TJD, JAP; Validation: PL, JAP; Visualization: PL, JAP; Writing – Original Draft Preparation: PL, JAP; Writing – Review & Editing: PL, KH, LM-G, ALZ, MSMB, TJD, JAP.

