## Supplementary Materials for "Exploring differences across pangenome-graph representations using *Escherichia coli* O157:H7 as a model"

### Supplementary Results

#### S1. Benchmark dataset description

To compare the pangenome representations produced by the selected tools, we compiled a dataset of *E. coli* O157:H7 complete genomes from RefSeq for which matching Illumina short-read data were available in the SRA database (see Methods). This serotype was chosen not only due to its public health relevance as a major foodborne pathogen, but also because its repeat-rich genome poses substantial challenges for assembly[1], providing a stringent test case for graph-construction methods.

Among all *E. coli* complete genomes in RefSeq (n = 2,980), 259 (8.69%) were annotated as O157:H7. Of these, 179 had corresponding Illumina datasets in SRA. After applying quality-control filters, 175 genomes were retained for downstream analyses. Accession numbers, SRA identifiers, and additional metadata are provided in Supplementary Table S2.

The phylogenetic composition of the dataset was strongly dominated by sequence type (ST) 11, which accounted for 172 isolates (98.3%) (Supplementary Figure S1A and S1B). The remaining isolates belonged to ST5560 and ST5516, both single-locus variants of ST11, and ST7816, a double-locus variant.

The fraction of repeated k-mers was calculated as the proportion of all k-mers in each genome that occurred more than once, providing a genome-wide measure of repetitive sequence content (Supplementary Figure S1C). O157:H7 isolates showed substantially higher repeat fractions (median = 0.0795; IQR = 0.0756–0.0884) compared with all other serotypes (median = 0.0384; IQR = 0.0309–0.0490). This difference was statistically significant based on a Wilcoxon rank-sum test (W = 665,298; two-sided, $p < 1\times10^{-16}$, $n_{O157:H7}=259$ , $n_{Other}=2,721$).

The O157:H7 serotype was highly enriched for Shiga toxin-producing *E. coli* (STEC)[2–4]. *In silico* virulence profiling showed that 168 isolates (96%) harbored at least one complete *stx* operon, composed of A and B subunits (Supplementary Table S2). Dual *stx* carriage was common: 87 isolates (49.1%) encoded two complete *stx* sets, with *stx1a-stx2a* (n = 35), *stx2a-stx2c* (n = 30), and *stx1a-stx2c* (n = 17) as the most frequent combinations (Supplementary Figure S1D). A further 76 isolates (43.4%) carried a single *stx* operon, most commonly *stx2c* (n = 45) or *stx2a* (n = 24). Rare configurations included three *stx* variants in four isolates and four variants in one isolate.

Because plasmids represent a major source of accessory genome variation and are frequently enriched in repetitive and mobile elements that challenge short-read assembly and graph construction, we analyzed the plasmidome of the benchmark dataset to contextualize downstream differences in pangenome-graph representations. A total of 284 plasmids were found in 173 genomes included in the benchmark, with only two genomes not carrying plasmids. Single-plasmid profiles were the most common (n = 81 isolates), followed by dual-plasmid profiles (n = 76). A smaller number of isolates carried three (n = 11), and four (n = 5) plasmid clusters. Plasmid backbones were assigned to clusters using *mge-cluster* (v1.1) coupled to the existing *E. coli* plasmid backbone database (Supplementary Figure S1E), as described in the methods section. Each isolate was assigned a plasmid profile defined by the set of clusters detected in its genome (Figure 1B). Of the 39 total distinct profiles, a small number of profiles dominated the dataset, with isolates carrying only plasmid cluster 25 (n = 47), only cluster 26 (n = 31), the combination 25;12 (n = 21), and 25;20 (n = 15) being the most prevalent. All remaining plasmid profiles occurred at substantially lower frequencies.

#### S2. Assembly-status distribution and detailed performance of *stx* loci recovery

The ground-truth burden comprised 266 *stx* operons (266 *stxA* and 266 *stxB* subunits). Assembly status was unevenly distributed between subunits. For *stxA*, 183/266 loci were Complete, 69/266 Fragmented, and 14/266 Collapsed. For *stxB*, 180/266 were Complete, 66/266 Fragmented, and 20/266 Collapsed (Supplementary Table S8).

Assembly artefacts were strongly structured by toxin background. Collapsed loci were predominantly observed in profiles consistent with duplicated or highly similar alleles. The most frequent collapsed configurations included "*stx1a; stx2a; stx2a*" (6 collapsed subunits each for *stxA* and *stxB*), "*stx2a; stx2a*" (6 collapsed *stxB* subunits), and "*stx1a; stx1a; stx2a; stx2a*" or "*stx2c; stx2c*" (4 collapsed subunits each). In contrast, Fragmented loci were enriched in mixed-variant backgrounds, particularly "*stx2a; stx2c*," which accounted for 60 fragmented *stxA* and 56 fragmented *stxB* subunits. Other multi-variant combinations were rare. These distributions confirm that both collapse and fragmentation preferentially affect isolates carrying multiple toxin variants and/or repeated alleles.

##### *a) Detailed performance for stxA*

For *stxA*, Panaroo and PPanGGOLiN maintained precision of 1.00 across all assembly-status strata. However, recall remained limited in collapse-prone and fragmented backgrounds. For Collapsed *stxA*, recall remained at 0.571 even in the fully complete dataset. For Fragmented loci, recall plateaued at 0.522 (Panaroo) and 0.507 (PPanGGOLiN) in the 100% complete condition.

ggCaller achieved substantially higher recall in Complete and Collapsed strata once ≥50% complete genomes were included, approaching 1.00. At 100% complete genomes, precision declined to 0.433 for Complete loci, 0.670 for Fragmented loci, and 0.778 for Collapsed loci, reflecting increased false-positive calls.

##### *b) Detailed performance for stxB*

For *stxB*, precision was near-maximal across tools and strata, with ggCaller showing a decrease to ~0.90 restricted to Complete loci when ≥50% complete genomes were present. Recall patterns mirrored those observed for *stxA*, with PPanGGOLiN reaching ~0.52 recall even in the fully complete dataset.

For Collapsed *stxB* loci, all pangenome methods remained below the Bakta baseline once datasets contained >50% complete genomes, with recall values remaining around ~0.6. These results indicate persistent under-recovery of collapsed toxin copies despite increasing availability of complete assemblies.

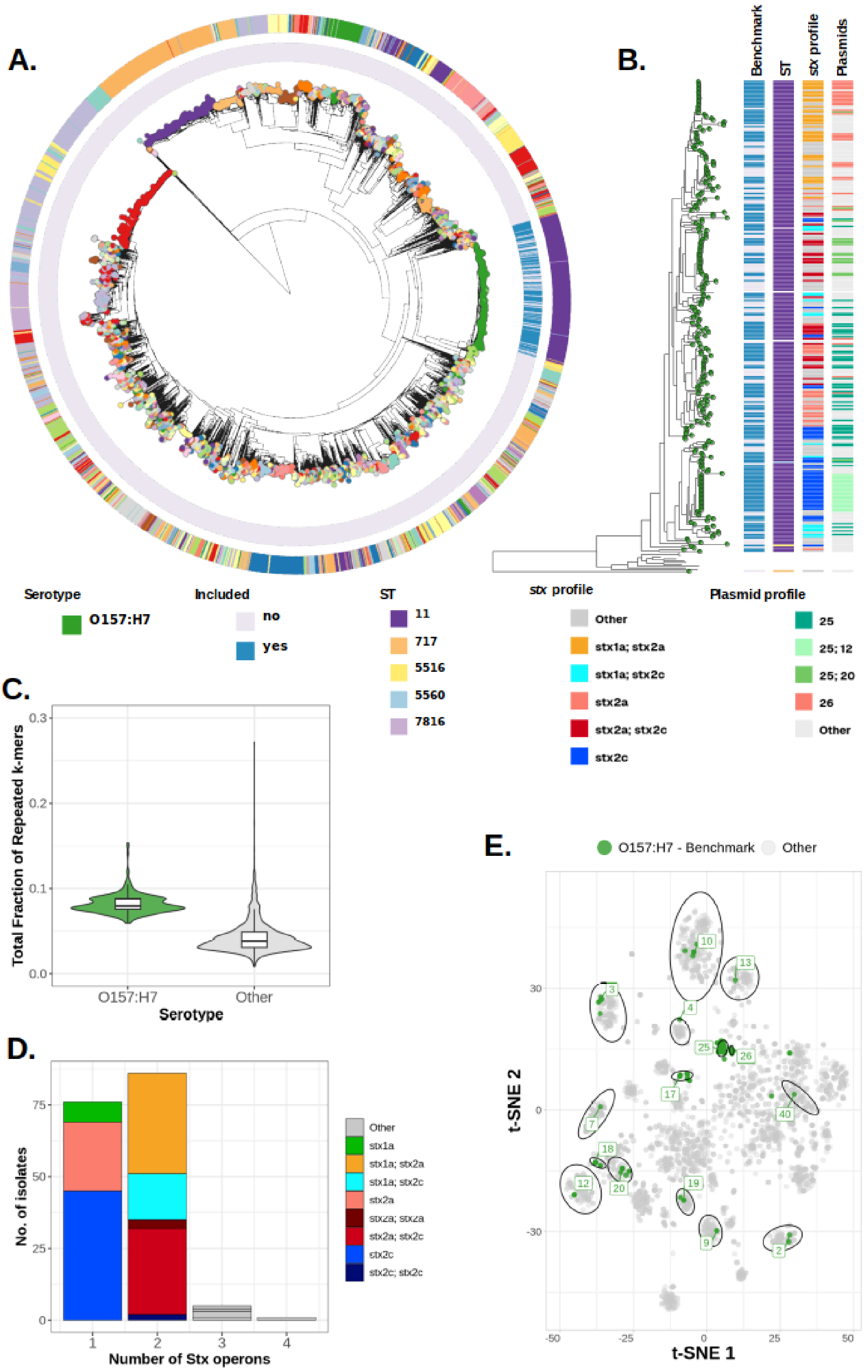


**Supplementary Figure S1. (A)** Neighbor-joining phylogeny of complete *E. coli* genomes from RefSeq (n = 2,980), with O157:H7 highlighted (green). Tracks indicate benchmark inclusion and MLST (O157-associated types shown). **(B)** Expanded phylogeny of benchmark O157:H7 isolates (n = 183) annotated with MLST, stx genotype, and plasmid profile (backbone clusters assigned using mge-cluster). **(C)** Genome-wide repeat content (fraction of k-mers occurring >1×) comparing O157:H7 and non-O157:H7 genomes; boxplots show median and IQR. **(D)** Distribution of complete stx operon counts (1–4) with variant combinations. **(E)** t-SNE projection of plasmid backbone clusters; benchmark plasmids are shown in green and reference plasmids in grey, with ellipses indicating clusters present in the benchmark.

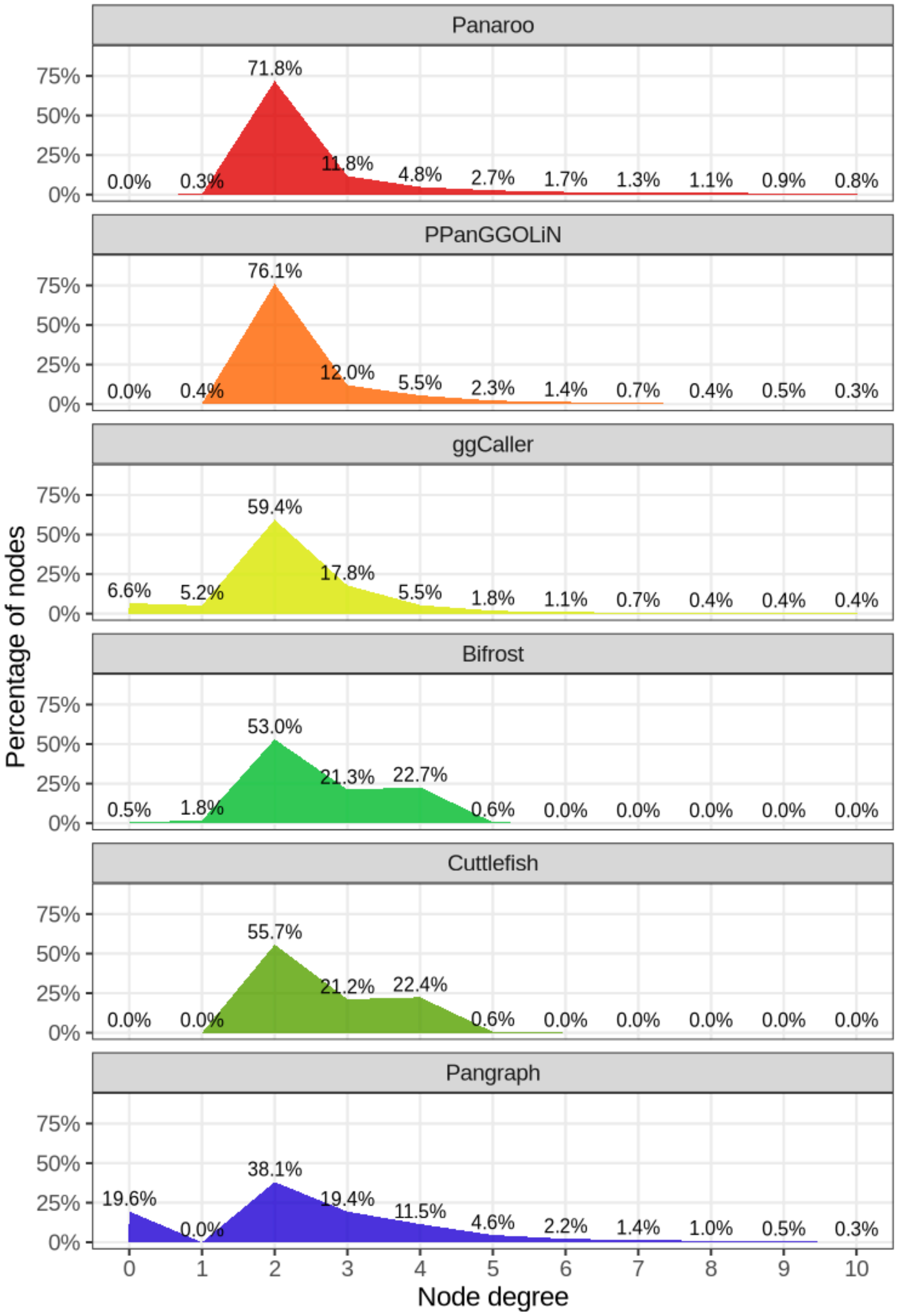


**Supplementary Figure S2.** These plots show the proportion of nodes at each node degree (0–10) per tool. Percent values are annotated at each degree.

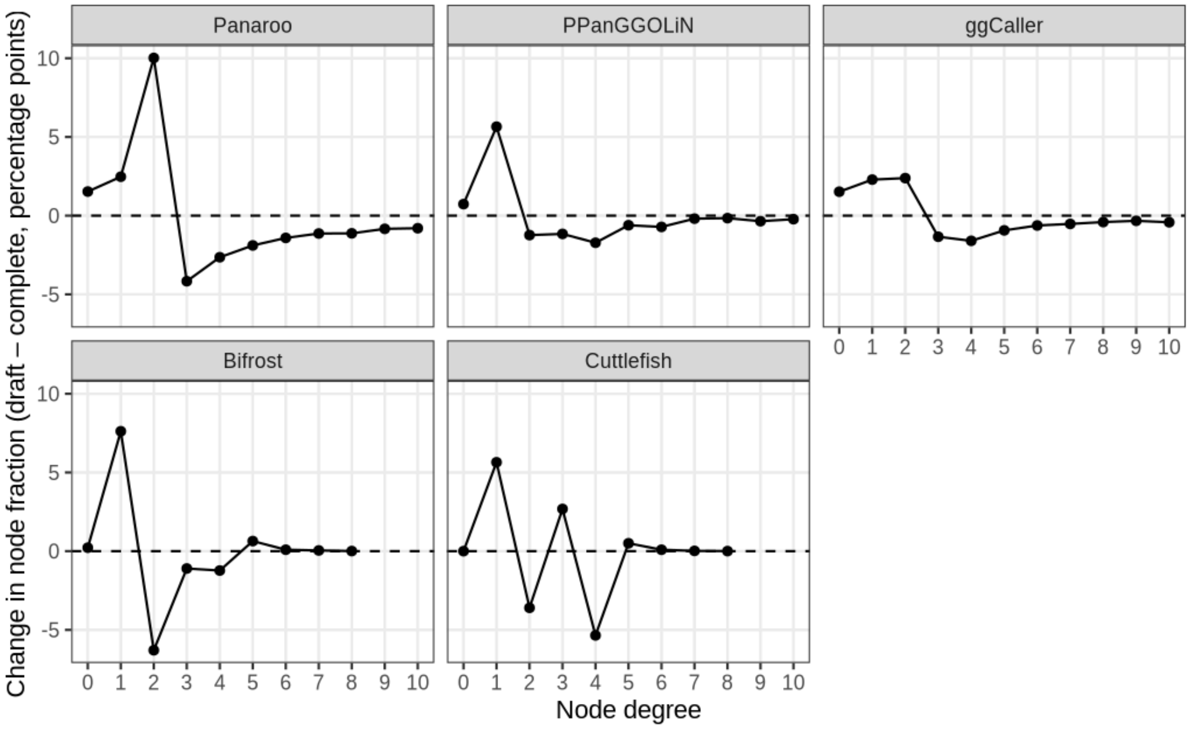


**Supplementary Figure S3.** For each method, the plot shows the change in the fraction of nodes at each node degree (x-axis; degrees 0–10) when moving from a graph built from 100% complete genomes to one built from 100% draft assemblies. Values are expressed as draft − complete in percentage points (y-axis); the dashed line marks no change. Positive values indicate enrichment of a given degree class under fragmented input, whereas negative values indicate depletion, summarizing how fragmentation shifts graph mass from higher-degree junctions toward lower-degree (0–2) nodes in a tool-specific manner.

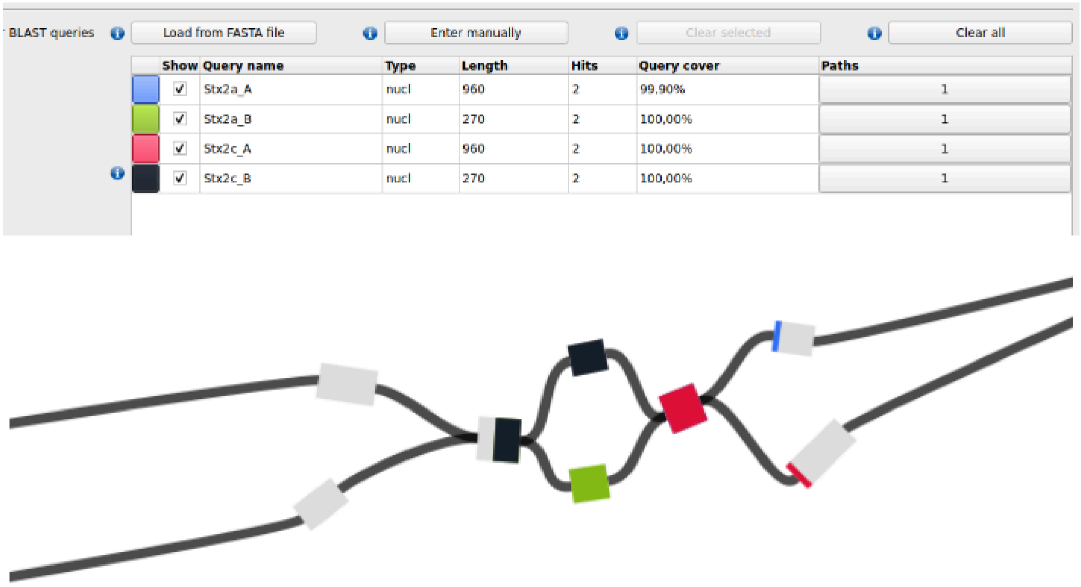


**Supplementary Figure S4.** Draft assembly graph section of genome visualized with Bandage. Each node in the graph represents a contig. Colored nodes (fragments) depict BLASTN hits against a *stx*-gene database. Blue fragments are *stx2a* subunit A; Green fragments are *stx2a* subunit B; Red fragments are *stx2c* subunit A; Black fragments are *stx2c* subunit B.

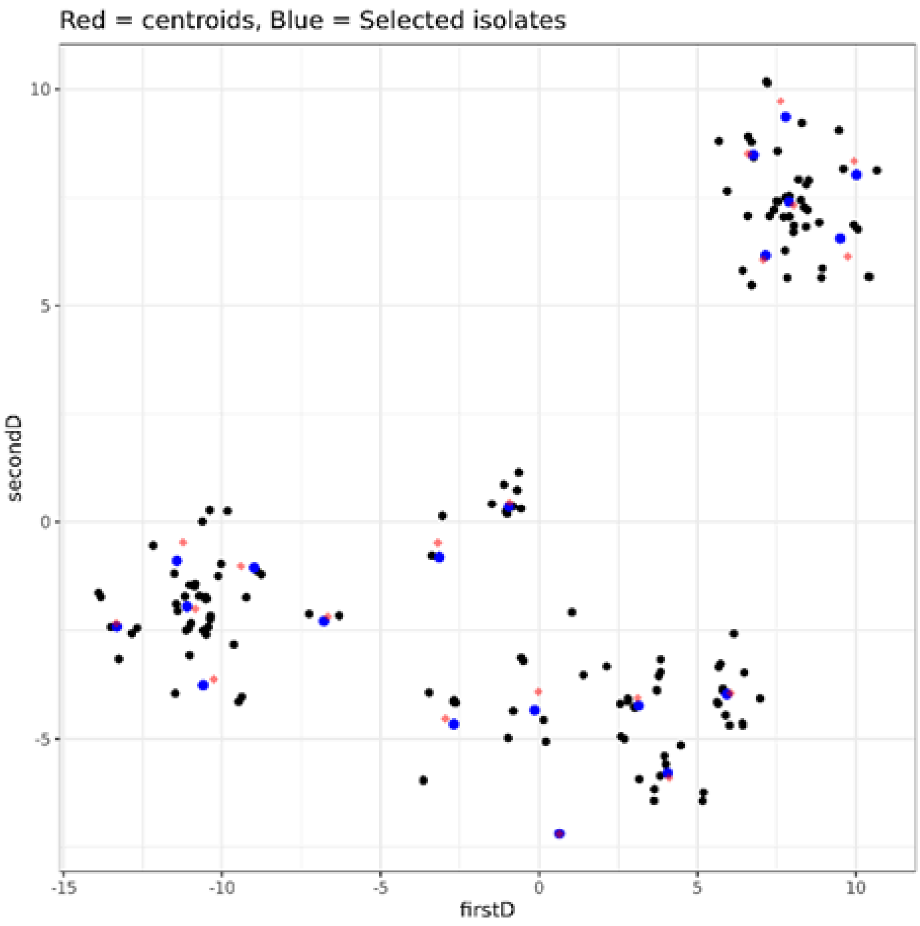


**Supplementary Figure S5.** Sample scatter plot for selecting 20 samples. Red dots represent the centers of the sample clusters, blue dots represent the selected samples.

### Supplementary Methods

#### Tool selection criteria

From the fourteen tools we found publicly available to build pangenome graphs (Supplementary Table S1), we selected six for downstream comparison: Panaroo (COG), PPanGGOLiN (COG), ggCaller (ccDBG + COG), Bifrost (ccDBG), Cuttlefish2 (ccDBG), and Pangraph (MSA). Rather than to provide an exhaustive comparison, we aimed to cover a set of tools that (i) collectively represent all major methodological paradigms for pangenome-graph construction and (ii) reflect the best-performing or most widely adopted implementations within each category. Where multiple tools existed within the same methodological class, selection was guided by prior benchmarking studies or, if unavailable, evidence of broad use within the microbial genomics community.

Roary was excluded due to repeated reports of inferior performance relative to Panaroo and ggCaller in terms of clustering accuracy and robustness to annotation errors[5,6] and because it is no longer actively maintained. Although Panaroo has a higher citation count than PPanGGOLiN, we included both tools because they implement fundamentally different strategies for modelling core and accessory genome structure[7] and, to date, no independent study has compared their performance.

For unitig-based methods, Bifrost and Cuttlefish2 were selected as the most widely used and best-supported ccDBG construction tools, as evidenced by citation counts (Supplementary Table S1) and extensive adoption both in microbial genomics workflows and as foundational components for downstream software[6,8,9]. Although GGCAT has been benchmarked against Bifrost and Cuttlefish2 and shown to achieve superior computational efficiency, these evaluations focused exclusively on runtime and memory usage rather than on the accuracy or biological representativeness of the compressed graph[10]. For this reason, we did not consider GGCAT as a replacement for Bifrost or Cuttlefish2 in the present comparison. Finally, we excluded mdbg because the tool requires low error long-read assemblies, and therefore falls outside the methodological scope of this benchmark.

ggCaller was included as the only method explicitly designed to integrate unitig calling with COG-based functional annotation, providing a hybrid representation of gene families and their genomic context. PanTools offers partially similar functionality, but its development and adoption have largely centred on eukaryotic pangenomes and large, complex genomes[11], making it less suitable for our microbial-focused evaluation.

Similarly, Pangraph was selected as the sole representative of the MSA-based paradigm, as it is the only tool in this category explicitly developed for microbial genomes and optimized for microbial-scale datasets[12].
